# A reaction-diffusion model captures the essence of liquid-liquid phase separation

**DOI:** 10.1101/2024.04.29.591603

**Authors:** Nayana Mukherjee, Abdul Wasim, Jagannath Mondal, Pushpita Ghosh

## Abstract

In this work we propose the formulation of a continuum model for liquid-liquid phase separation (LLPS) using reaction diffusion framework. We consider a well mixed liquid consisting of three phases, the dense droplet phase, the dilute phase and the remaining part to be solvent phase. As a key feature, the model includes both the spatial and temporal aspects and a free energy functional between dense and dilute phase that is physically motivated by reference molecular simulation. The exhaustive numerical simulations of model captures the dynamical formation of droplets and existence of LLPS. As the time progresses, simulation reveal that smaller droplets gradually vanish, and a single droplet undergoes continuous growth until it reaches a stable size. The model predicts that that extent of diffusivity of dense and dilute phase as well as their mutual interaction would modulate the kinetics of droplet formation. Finally we show that introduction of fluctuation in the model accelerate the phase separation process.

## I. INTRODUCTION

Liquid-liquid phase separation (LLPS) is a phenomenon of significant interest, characterized by the spontaneous partitioning of a uniform solution or mixture into two distinct phases, resulting in regions with markedly different component concentrations. Prominent examples include the emergence of membrane-less organelles (MLOs) such as Cajal bodies [1] and the formation of in-vivo droplets by intrinsically disordered proteins (IDPs). While MLOs play essential roles, the in-vivo LLPS of IDPs has been shown to be highly pathogenic [2–4]. These LLPS events, observed both in vivo and in vitro, are significantly influenced by factors such as temperature, ionic strength, and the presence of crowding agents.

State-of-the-art experiments have greatly expanded our understanding of liquid-liquid phase separation (LLPS) processes and their intricacies [5–11]. However, a comprehensive physical grasp of LLPS remains a goal. To pursue this, a burgeoning approach involves simulating these phenomena using computationally derived agent-based models[12–19]. While molecular models strive to provide detailed insights, they are limited by system size and time scale constraints. Hence, a fast and accurate model capable of simulating experimental time and length scales, while capturing essential aspects of phase separation, would be a significant leap forward. A continuum model of LLPS aligns with the objectives which we aim to develope in this study.

Classical theories have endeavored to characterize nucleation, growth, and coarsening in liquid-liquid phase separation (LLPS), encompassing active and passive emulsions, binary mixtures, and droplet dynamics [20, 21]. Predominant approaches, rooted in classical continuum modeling, offer a comprehensive understanding of LLPS types observed in experiments and associated modeling techniques. These models typically formulate dynamical equations to elucidate droplet coarsening resulting from coalescence. Investigations have explored LLPS modeling using JMAK theory, the Ginzburg-Landau equation, and the Flory-Huggins equation, addressing various LLPS stages [22–24]. A recurrent challenge in model development is the selection of a suitable free energy functional to determine concentration phase spatial distributions at thermal equilibrium. Statistical approaches to free energy functionals for instabilities are explored in complex mixtures. In chemical reactions, phase segregation dynamics are observed in binary mixtures, employing the Ginzburg-Landau functional for the free energy term [25]. While the Cahn-Hilliard equation is widely used within the reaction-diffusion framework, the rationale for selecting a specific free energy functional remains unclear across literature [26–30].

The aforementioned discussion prompts an exploration into whether a simpler continuum-based approach can effectively model LLPS. A reaction-diffusion framework might help to achieve our goal since LLPS has both spatial and temporal aspects to be explored in detail. We aim to formulate the model such that it can capture all the important properties of LLPS. Apart from modeling the nucleation and aggregation processes of droplets, we need to include the free-energies of the different phases into the model. Instead of a using general free energy functional term, we can actually calculate the same from molecular dynamical simulations and give it a functional form which models interaction between droplet and dilute phases. With the help of analytical tools like linear stability analysis, we can get an idea of the parameter regimes for which the model will show proper LLPS and droplet formation in simulations. This type of coarsegrained modeling will help in bringing down the computation time comparatively but will help to understand the broad aspects of LLPS completely from the exhaustive numerical simulations. We dig deep into the modeling approach in the next section and the associated results and discussions are presented in the subsequent chapters.

## II. MODEL AND METHODS

### A. The Model

To formulate a comprehensive model for Liquid-Liquid Phase Separation (LLPS), we begin with the fundamental assumption of a system comprising of at least two components with distinct densities. One of these components represents the dense phase responsible for droplet formation, while the other signifies the dilute phase. The subsequent step involves modeling the intricate processes of nucleation and aggregation of droplets, culminating in the manifestation of phase separation.

In our approach, we adopt a reaction-diffusion framework to elucidate the intricate process of droplet formation within a confined spatial domain, denoted as [0, *L*] × [0, *L*]. Herein, the variables *N* (*x, y, t*), *P* (*x, y, t*), and *W* (*x, y, t*) are assigned to represent the droplet component, dilute phase, and the more dilute or water phase, respectively (refer to Fig.1). With these considerations in mind, we have devised a model presented as a system of partial differential equations, commonly referred to as a reaction-diffusion system. The model is characterized by non-zero initial conditions and no-flux boundary conditions.The governing equations take the following form:

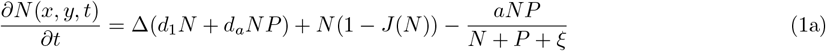

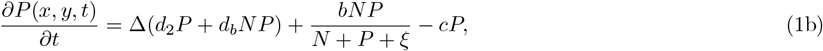

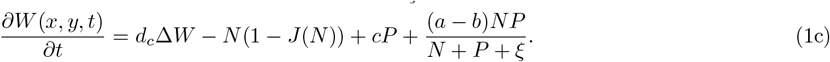

Where

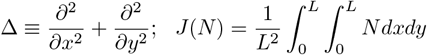

**FIG. 1:**
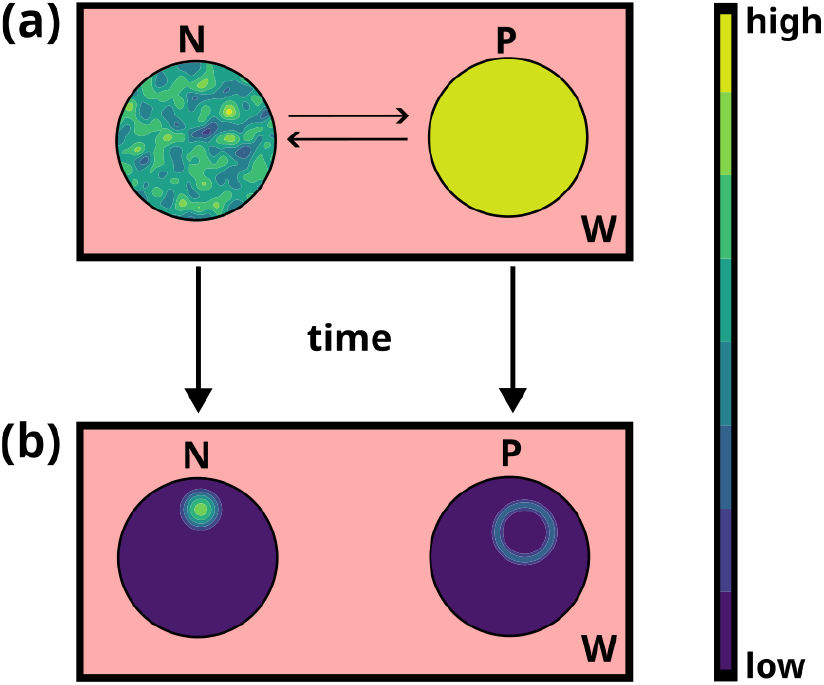
This is a schematic representation for the mathematical model given in (1): N is the droplet phase, *P* is the dilute phase and *W* is the remaining part of the liquid: (a) This represents the initial well mixed state where all the components are equally distributed over the considered bounded domain; (b) This depicts the later stage where the stable droplet *N* has formed and cannot be converted to dilute phase *P* anymore.

The left hand side part denotes the change of the different phases with respect to time whereas the right hand side of Eq. 1 takes care of the diffusion and the reaction terms specific to each component. We describe in detail each of the terms and constants in the following paragraphs.

#### The Interaction among dense phases (N -N interaction term)

The term *N* (1 −*J*(*N*)) represents the interaction between *N* components, influencing the growth and eventual nucleation of droplets. Ideally, this interaction term is intended to mimic logistic growth, expressed as *N* (1 −*N*). In this formulation, we also incorporate the modeling of aggregation phenomenon. We introduce *J*(*N*) instead of *N*, to take care of the aggregation. It is the global average term, where the average of the *N* phase concentration is considered over the whole spatial domain. This term also takes care of the interaction of *N* components, not only locally around the considered spatial point but interactions anywhere across the entire domain [31].

#### The interaction between dense and dilute phases (N -P, P -P interaction terms)

The functional form for the *N* -*P* interaction is 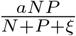, which is derived from concurrent molecular dynamics simulations aimed at investigating the aggregation phenomenon in the *α*-Synuclein protein. We consider a distribution of *α*-Synuclein in various stages of aggregation, where the mole fraction 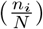 of each cluster or monomer is denoted as *ϕ*_*i*_ (details given in SI). It represents the part of concentration that diminishes from the *N* phase and contributes to the dilute phase *P*. This explains its inclusion in the growth term of *P*, since it helps in the growth of *P* phase, albeit scaled by a conversion coefficient 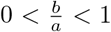. The *N* -*P* interaction term, formulated as 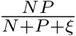, is contingent upon the overall concentration of *N* and *P* across the entire domain, ensuring normalization. The constant *ξ* serves as a saturation coefficient beyond which *P* converts to *N*, playing a crucial role in droplet formation. The positive constants *a* and *b* signify the rate of decay of the *N* phase and the rate of growth of the *P* phase, respectively, with the condition *a > b*. Additionally, the *P* -*P* interaction is assumed to adopt a linear form, represented as −*cP*, symbolizing the transfer of the dilute phase into the droplet or water phase.

#### Diffusion and cross-diffusion of three phases

The positive constants *d*_1_, *d*_2_, and *d*_3_ represent the self-diffusion coefficients, serving as rates of diffusion for the *N, P*, and *W* components, respectively. The terms *d*_1_Δ*N, d*_2_Δ*P* and *d*_3_Δ*W* account for the random diffusion of the droplet, dilute, and water phases in the given mixture. Additionally, we incorporate cross-diffusion terms into the model to capture the influence of the *N* component on the movement of the *P* component, and vice versa. This implies that the movement of one component is influenced by the presence, absence, abundance, or scarcity of the other component and vice versa. The cross-diffusion coefficients *d*_*a*_ and *d*_*b*_ are both positive. An essential motivation for introducing cross-diffusion terms in our model is to avoid a direct correlation between the components *N* and *P*, preventing high-density areas of *N* from aligning with high-density areas of *P* and vice versa. This helps prevent undesired patterns in the system. Coarse grained molecular dynamics simulations using a modified Martini 3, adapted for *α −synuclein* [12], were performed on an aggregate (14 mer) and a single chain of *α*-synuclein for determination of their mean squared displacements (MSDs) (details given in SI). We got the ratio of *D*_*monomer*_ to *D*_*oligomer*_ to be 12.52 according to the simulations. So we use the ratio for the droplet to dilute phase to be 1*/*10. It is observed that the ratios 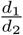 or 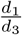 are on the order of 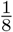. Therefore, we choose the values of *d*_1_, *d*_2_, and *d*_3_ in a consistent manner, maintaining this ratio. Also we have experimental results which say that the diffusion of droplet or dense phase (*N*) is restricted [6, 32].

#### Solvent component or more dilute phase

Given the consideration of two liquids in the mixture, it is essential to maintain a constant concentration of the mixture. To achieve this, we define *W* (*x, y, t*) to encompass the remaining elements of the mixture as a background. In the absence of spatial dependence, the PDE system transform to ODE system, the term *J*(*N*) also transforms into *N*, leading to 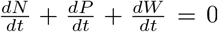. Integrating this expression reveals that *N* + *P* + *W* is a constant, signifying the maintenance of a constant concentration in our model.

## III. RESULTS AND DISCUSSION

### A. Simulation results exhibit the formation of droplets and existence of LLPS

This is a representative case where LLPS and droplet formation can be affirmed via numerical simulations. We proceed with numerical simulations of the spatially extended system, integrating system (1) in two dimensions. The explicit Euler method is employed for numerical simulations, with a discretization of both space and time. We utilize a finite system size with dimensions *L*_*x*_ = *L*_*y*_ = 200, employing a grid size of Δ*x* = Δ*y* = 1.0, and a time interval *dt* = 0.0001 for accuracy. Throughout our study, we maintain a zero-flux boundary condition. The initial state of the system, initialized at each mesh point in a 200×200 array, is perturbed by introducing ±1% random noise. This intentional perturbation breaks the initial spatial symmetry and ensures a more realistic representation of the system’s evolution. To ensure the stability and persistence of stable heterogeneous droplet formation, we extend the numerical simulations over an elongated time period. This comprehensive approach allows us to observe and analyze the dynamics and patterns that emerge in the system under various conditions.

#### Droplet formation and their evolution

Figure 2 visually depicts the formation of droplets through liquid-liquid phase separation (LLPS). In the sequence of figures (a-d), it is evident that the chosen parameter set with *a* = 1.9 leads to the emergence of droplets when an initial mixture of *N, P*, and *W* undergoes simulation using the complete nonlinear system (1). After a few hundred time steps, observable droplets begin to appear, exhibiting considerable size variations. As time progresses, smaller droplets diminish, while larger ones undergo substantial growth. After thousands of time steps, a single dominant droplet persists across the entire domain.

**FIG. 2:**
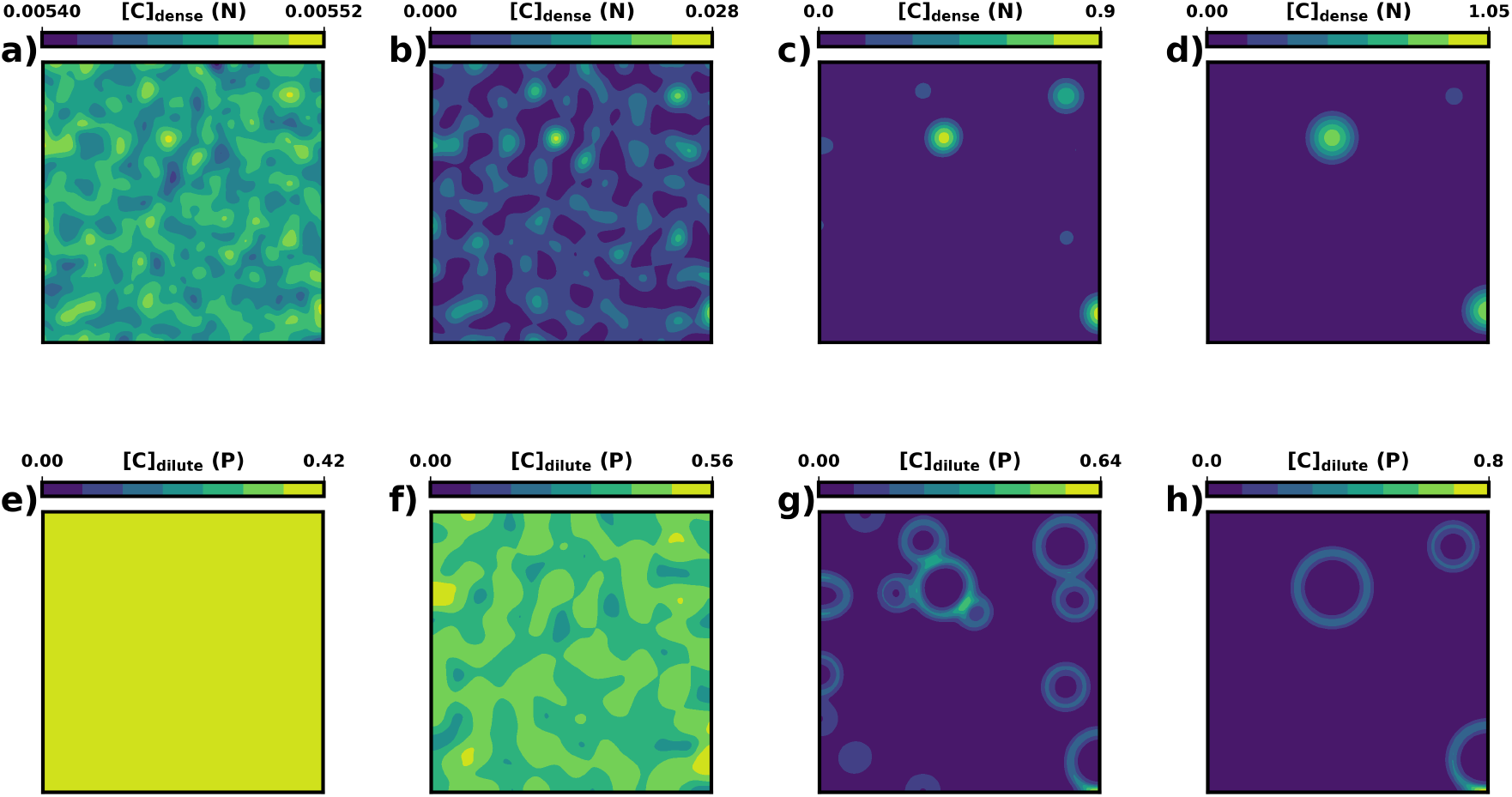
Evolution of droplets with advancement of time for parameter values *a* = 1.9, *b* = 1, *c* = 0.6, *ξ* = 0.001 *d*_1_ = 1, *d*_2_ = 10 = *d*_3_ = *d*_*a*_ = *d*_*b*_: upper panel is for droplet phase *N* and lower panel is for dilute phase *P* : (a,e) 100 time point; (b,f) 300 time point; (c,g) 1000 time point; (d,h) 3000 time point.

This observed phenomenon aligns with Ostwald ripening. Notably, before the final formation of a single droplet, the last two droplets, one with low concentration and another with high concentration, illustrate the characteristic behavior of Ostwald ripening. The smaller droplet gradually diminishes and ultimately disappears, leading to the continued growth of the larger droplet with higher concentration as time evolves [refer to Fig. 2(d)]. The aggregation term *J*(*N*) from the model (1) plays a crucial role in the aggregation of *N* components. Without the inclusion of *J*(*N*), the coalescence of droplets would not be observable. However, with the presence of *J*(*N*), we observe the aggregation of droplets into a single, larger droplet. The aggregation process can manifest in various ways, with Ostwald ripening being a common occurrence. In Ostwald ripening, smaller droplets diminish over time, while larger droplets grow, eventually forming the largest droplet in the system. A subsequent analysis follows the characterization of the above mentioned observation accompanying the variation of droplet sizes leading to a bigger one.

To characterize the emerging droplets observed in nu-merical simulations, we employ a method that involves tracking the evolution of droplet size over time. As the simulation progresses, smaller droplets gradually vanish, and a single droplet undergoes continuous growth until it reaches a stable size. At this juncture, the droplet’s size stabilizes, and its radius is calculated. For the analysis, we utilize the OpenCV Python package, leveraging its implementation of the Hough Circle transform algorithm. This algorithm enables the detection of circles, allowing us to estimate their radii. When the droplets reach a significant size and the liquid-liquid phase separation (LLPS) process is stable, we apply this technique to detect the circles representing the circumferences of the droplets [refer to Fig. 3(a)]. Subsequently, we calculate the change in radius of the dominant droplet over time. Observations from these analyses reveal a gradual decrease in the number of droplets over time, eventually converging to a single stable droplet, as illustrated in Figure 3(b). This analysis provides valuable insights into the evolution and stability of droplets during the LLPS process. Moreover we also observe the radius of the largest droplet keeps increasing until it reaches a maxima after a timepoint (Fig. 3c). We now try to vary the parameter values to see how far can the LLPS be seen using numerical simulation for which we need some analytical help.

**FIG. 3:**
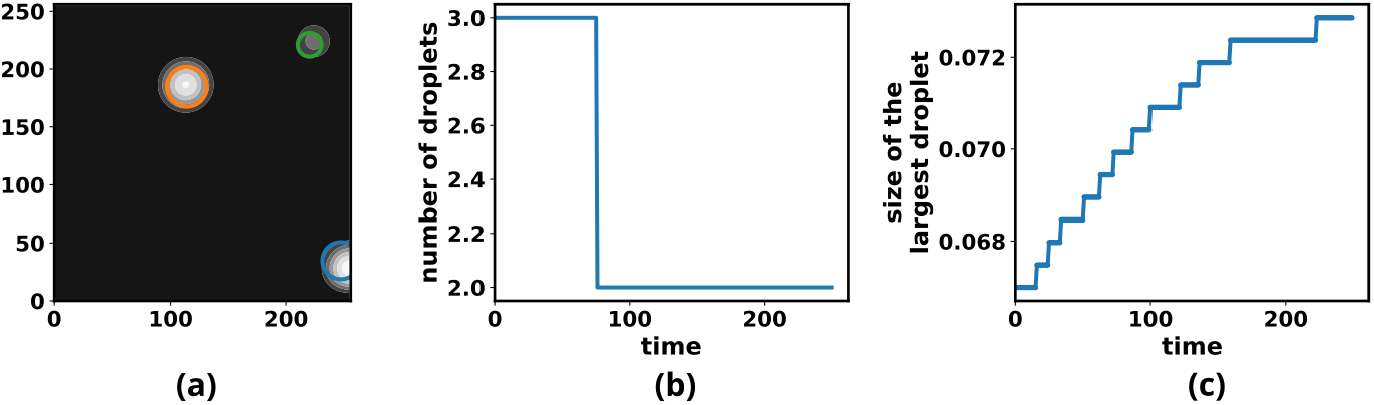
(a) The radii detected from circles detected from droplet phase (*N*) and dilute phase (*P*) for the parameter values *a* = 1.9, *b* = 1, *c* = 0.6, *ξ* = 0.001 *d*_1_ = 1, *d*_2_ = 10 = *d*_3_ = *d*_*a*_ = *d*_*b*_; (b) The number of the droplets coming down from 3 to 2 for the parameter values *a* = 1.9, *b* = 1, *c* = 0.6, *ξ* = 0.001 *d*_1_ = 1, *d*_2_ = 10 = *d*_3_ = *d*_*a*_ = *d*_*b*_ versus time; (c) The maximum size of droplets versus time; the corresponding radius of the stable droplet starts increasing but eventually stabilizes to a fixed one.

### B. Determination of parameter space for droplet formation via stability analysis

To comprehend the dynamics of model (1), we employ linear stability analysis, and the detailed procedure and associated Fig. 4. A two parameter bifurcation diagram in the *a*-*d*_2_ plane as shown in Figure 4, demonstrates that as *a* increases-indicating an intensified *N* -*P* interaction - the critical values of *d*_2_ also increase. The parameter values are taken as *b* = 1, *c* = 0.6, *ξ* = 0.001, *d*_1_ = 1.0, *d*_3_ = 10.0 = *d*_*a*_ = *d*_*b*_ and the *N* -*P* interaction parameter *a* is ranged from 1.5 - 2.1. The critical values delineate the boundary below which droplets do not form and above which LLPS is observable. Notably, when considering the diffusion of the dilute phase beyond the critical value (illustrated by the blue curve), diffusion driven patterns is evident. For higher values of diffusion one can see proper droplet formation as shown in Fig. 4. Conversely, the black dashed line represents the Hopf bifurcation curve, with the threshold for Hopf bifurcation denoted as *a*_*H*_ = 1.95. For values of *a < a*_*H*_, the homogeneous steady state is stable against temporal perturbations and vice versa. However, the presence of LLPS is observable for values of *d*_2_ surpassing the critical threshold. Our examination, based on a selected set of parameter values, reveals that within a specific interval in the range of (0, 1), the real part of eigenvalues *Re*(*λ*) becomes positive. The resulted plot is shown in Figure 2(c) from SI with respect to the wave number *k*. This marks the onset of diffusive instability, and the presence of liquid-liquid phase separation (LLPS) can be observed through numerical simulations.

**FIG. 4:**
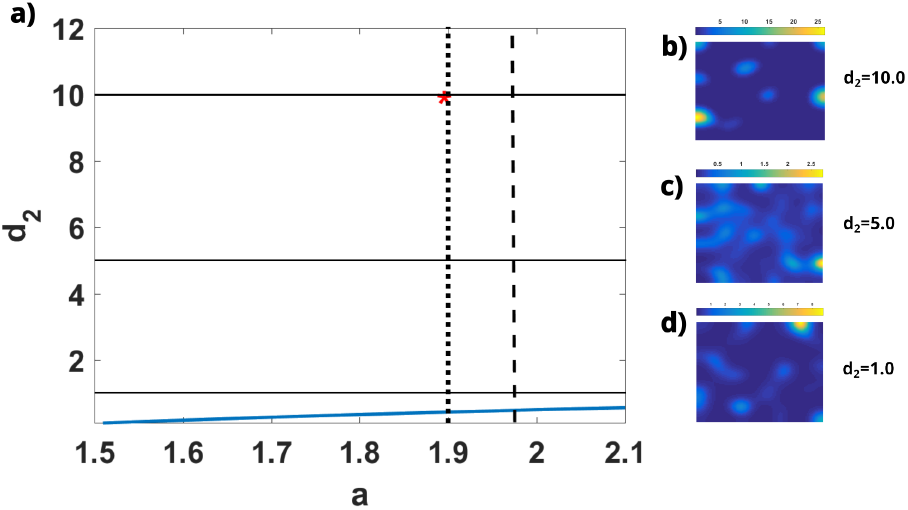
Two parameter bifurcation diagram in *a* - *d*_2_ plane for parameter values: *b* = 1.0, *c* = 0.6, *ξ* = 0.001, *d*_1_ = 1.0, *d*_3_ = 10 = *d*_*a*_ = *d*_*b*_, black dashed straight line represents the Hopf bifurcation curve, blue curve represents the diffusion induced instability curve, red ‘*’ represents the parameter values *a* = 1.9, *d*_2_ = 10 which is used in numerical simulations; Plots of *N* phase for (b) *d*_2_ = 10; (c) *d*_2_ = 5; (d) *d*_2_ = 1.

#### Change of diffusivity constants affect droplet formation

Considering the diffusion and cross-diffusion parameters, we aim to examine their influence on the LLPS process. Utilizing a set of kinetic parameters with *a* = 1.9, *b* = 1, *c* = 0.6, and *ξ* = 0.001 as the baseline, we proceed to vary the parameters *d*_2_, *d*_3_, *d*_*a*_, and *d*_*b*_, while maintaining *d*_1_ = 1. Numerical simulations of the system using the parameter values as given in caption of Fig. 4, reveal that for lower values of *d*_2_ the solutions exhibit movement within the domain, forming small aggregates that subsequently break apart. This process continues, and although droplets do not form, the solutions move around in the domain, creating irregular wavelike structures but an aggregation definitely occurs (see Figs. 4(c) and (d)). But for higher values of *d*_2_ we see proper droplets to form as shown in Fig. 4(b).

Contrastingly, when diffusion parameter is considered to be much higher like 10, droplet formation occurs. Although, linear stability analysis and resulting dispersion curves as shown in Figure 3 in SI show positive eigenvalues for a fixed *d*_2_ = *d*_3_ = 0.1, 1.0 and 5.0 values and with varied cross-diffusion parameters, it requires a significantly high value of diffusion constants for the droplet formation. Also, the plots reveals that rate of aggregation and droplet formation increases with higher values of *d*_2_. This observation emphasizes the intricate interplay between the diffusion and cross-diffusion parameters in influencing the dynamics of liquid-liquid phase separation in the system.

#### Extent of N -P interaction modulates the kinetics of phase separation

In examining the impact of the interaction parameter *a* on droplet formation, we observe distinct changes in the dynamics, particularly as the degree of *N* -*P* interaction (variation of parameter ‘a’) is heightened. Analyzing the plots of *Re*(*λ*) for various values of *a* as demonstrated in Fig. 5(a), it is evident that with an increase in *a* the maximum growth rate increases. As *a* is elevated, the system (1) tends toward faster instability and the formation of droplets becomes more rapid, leading to an accelerated aggregation process. The parameter *a* accounts for the decay of the *N* phase to the *P* phase. However, as the density of the dilute phase increases and settles around the *N* droplet phase, the aggregation of droplets intensifies.

**FIG. 5:**
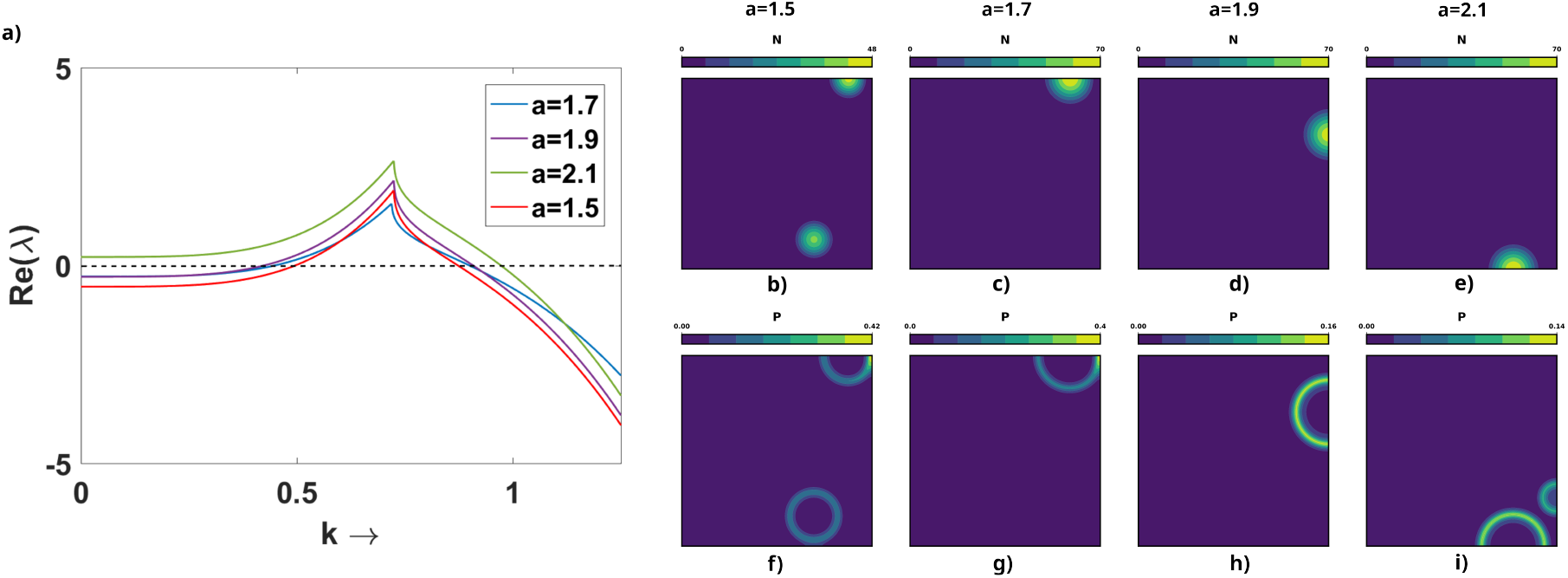
(a) Plot of the *Re*(*λ*) from the linear stability analysis for varying values of *a* and *b* = 1, *c* = 0.6, *ξ* = 0.001 *d*_1_ = 1, *d*_2_ = 10 = *d*_3_ = *d*_*a*_ = *d*_*b*_; Evolution of droplets after one lac timepoint: (b),(f) *a* = 1.5; (c),(g) *a* = 1.7; (d),(h) *a* = 1.9; (e),(i) *a* = 2.1.

**FIG. 6:**
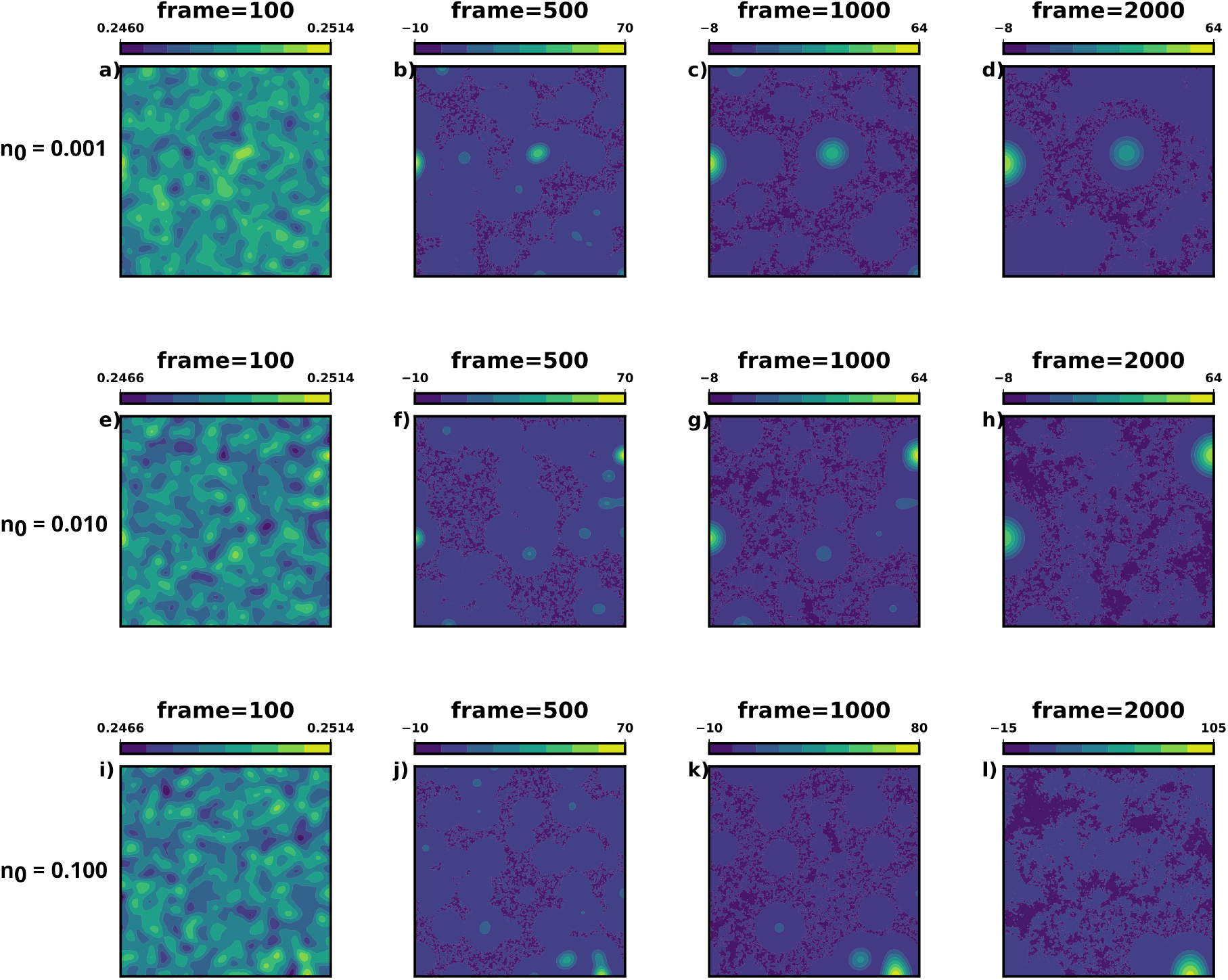
(a)-(d): Formation of droplets for *n*_0_ = 0.001. (e)-(h): Formation of droplets for *n*_0_ = 0.010. (i)-(l): Formation of droplets for *n*_0_ = 0.100.

The corresponding numerically simulated patterns are illustrated in Fig. 5(b)-(i). The entire upper row in Figs. 5 (b), (c), (d), and (e) depicts the evolution of the droplet phase N. By varying the values of the N -P interaction parameter a from 1.5 to 2.1 in simulations of the model (1), we observe a notable acceleration in the rate of aggregation. The corresponding lower row of Figs. 5 (f), (g), (h) and (i) displays the dilute phase P, which aggregates around the droplets. The density of the dilute phase notably increases with an elevation in the value of a. This behavior aligns with the faster aggregation observed in the droplet phase. Numerical simulations affirm that aggregation accelerates as a increases, while analytical insights indicate that the system becomes more unstable with an augmentation of a. This comprehensive analysis underscores the intricate relationship between the interaction parameter and the dynamic behavior of droplet formation in the considered system.

### C. Noise induced phase separation observed

Since fluctuation is ubiquitous, next we incoporate an aditive white noise into the droplet phase of the spatially-extended system as

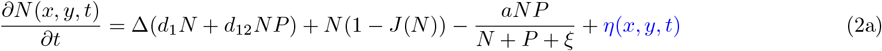

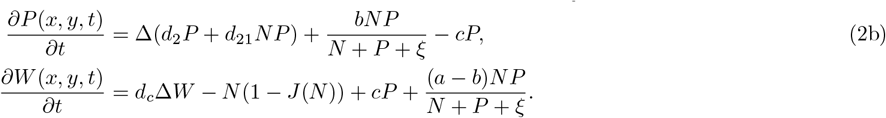

Where

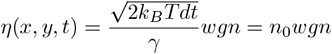

where “wgn” refers to white Gaussian noise. Time taken is one lac time units with *dt* = 0.0001. Other parameters are *a* = 1.9.*b* = 1, *c* = 0.6, *d*_*N*_ = 1, *d*_*P*_ = 10, *d*_*a*_ = 1, *d*_*b*_ = 10, *d*_*W*_ = 10. We consider the noise term changes at every time step, it is taken within the time loop as per numerical simulations. We consider the constant values for Δ_*t*_, *γ*, and we take values of *n*_0_ from 10^*−*3^, 10^*−*2^ and 10^*−*1^ to see the effect of noise is actually the effect of increasing temperature since the term depends on the value of *k*_*B*_*T*. As *n*_0_ is increased from 0.0001(top one) through 0.001(second), 0.01(third) to 0.1(bottom) we see that the droplets form more quickly. When the intensity is low, the time taken to form the droplets is much higher than when the intensity of noise is much higher, i.e. the droplets form more quickly then. Also as the strength of noise increases, we see it takes less time for the droplets to converge to single droplet. So, we can conclude that this is more of a noise induced phase separation. This is also to be mentioned that even if we use one of the noise terms, out of *ξ*_1_ and *ξ*_2_, then also we observe similar results.

## IV. CONCLUDING REMARKS

This work depicts the formulation of a generic mathematical model using reaction-diffusion framework which when numerically simulated can capture liquid-liquid phase separation and droplet formation. All the terms considered represent the various interactions taking place in LLPS. Apart from the droplet and dilute phase, we have considered the water phase which takes care of the remaining part of the liquid, such that the concentration of the liquid is constant. The relevant analytical and numerical tools has helped us to find appropriate parameter space where droplets can be seen and the simulation results have proved them right. We can see small droplets forming and eventually aggregating into a single droplet which is the case seen in Ostwald ripening. The relevant numerical techniques for characterization of the droplets have shown the number of droplets decreasing with and correspondingly the radius of the bigger droplet increasing until it becomes the single stable droplet. The change of the interaction parameter between droplet and dilute phase heavily impacts the speed of the droplet formation. Also when the diffusion and cross-diffusion constants are of very high range, we see proper droplet formation. Otherwise we can see aggregation of the solutions but not droplets. Also introduction of white Gaussian noise into the system leads to noise-induced phase separation, which is quite an interesting result to follow. In conclusion this is our attempt to provide the simplistic generic mathematical model for LLPS. It cannot capture the more complicated phenomena associated with LLPS like droplets within droplets seen in active emulsions. But if we use necessary modifications, we might be able to model such situations with reaction-diffusion framework. We look forward to use this idea to capture more complex results of LLPS.

## ACKNOWLEDGMENTS

We acknowledge support of the Department of Atomic Energy, Government of India, under Project Identification No. RTI 4007. JM acknowledges Core Research grants provided by the Department of Science and Technology (DST) of India (CRG/2023/001426). PG acknowledges Start-up Research Grant (SRG/2022/000043) by DST. India

## References

[1] T. Hirose, K. Ninomiya, S. Nakagawa, and T. Yamazaki, A guide to membraneless organelles and their various roles in gene regulation, Nature Reviews Molecular Cell Biology 24, 288 (2023).

[2] D. L. Price, S. S. Sisodia, and S. E. Gandy, Amyloid beta amyloidosis in alzheimer’s disease, Current opinion in neurology 8, 268 (1995).

[3] L. Breydo, J. W. Wu, and V. N. Uversky, α-synuclein misfolding and parkinson’s disease, Biochimica et Biophysica Acta (BBA)-Molecular Basis of Disease 1822, 261 (2012).

[4] A. Agarwal and S. Mukhopadhyay, Prion protein biology through the lens of liquid-liquid phase separation, Journal of Molecular Biology 434, 167368 (2022).

[5] S. Wegmann, B. Eftekharzadeh, K. Tepper, K. M. Zoltowska, R. E. Bennett, S. Dujardin, P. R. Laskowski, MacKenzie, T. Kamath, C. Commins, et al., Tau protein liquid–liquid phase separation can initiate tau aggregation, The EMBO journal 37, e98049 (2018).

[6] S. Ray, N. Singh, R. Kumar, K. Patel, S. Pandey, D. Datta, J. Mahato, R. Panigrahi, A. Navalkar, S. Mehra, et al., α-synuclein aggregation nucleates through liquid–liquid phase separation, Nature chemistry 12, 705 (2020).

[7] S. Sawner, S. Ray, P. Yadav, S. Mukherjee, R. Panigrahi, M. Poudyal, K. Patel, D. Ghosh, E. Kummerant, Kumar, et al., Modulating α-synuclein liquid–liquid phase separation: Published as part of the biochemistry virtual special issue “protein condensates”, Biochemistry 60, 3676 (2021).

[8] Y. Lin, Y. Fichou, A. P. Longhini, L. C. Llanes, P. Yin, G. C. Bazan, K. S. Kosik, and S. Han, Liquid-liquid phase separation of tau driven by hydrophobic interaction facilitates fibrillization of tau, Journal of molecular biology 433, 166731 (2021).

[9] S. Mukherjee, A. Sakunthala, L. Gadhe, M. Poudyal, A. S. Sawner, P. Kadu, and S. K. Maji, Liquid-liquid phase separation of α-synuclein: a new mechanistic insight for α-synuclein aggregation associated with parkinson’s disease pathogenesis, Journal of molecular biology 435, 167713 (2023).

[10] S. Sudhakar, A. Manohar, and E. Mani, Liquid–liquid phase separation (llps)-driven fibrilization of amyloid-β protein, ACS Chemical Neuroscience 14, 3655 (2023).

[11] Z. Zhang, G. Huang, Z. Song, A. J. Gatch, and F. Ding, Amyloid aggregation and liquid–liquid phase separation from the perspective of phase transitions, The Journal of Physical Chemistry B 127, 6241 (2023).

[12] Wasim, S. Menon, and J. Mondal, Modulation of α-synuclein aggregation amid diverse environmental perturbation, bioRxiv, 2023 (2023).

[13] S. Mukherjee and L. V. Schäfer, Thermodynamic forces from protein and water govern condensate formation of an intrinsically disordered protein domain, Nature Communications 14, 5892 (2023).

[14] G. L. Dignon, W. Zheng, Y. C. Kim, R. B. Best, and J. Mittal, Sequence determinants of protein phase behavior from a coarse-grained model, PLoS computational biology 14, e1005941 (2018).

[15] G. Tesei and K. Lindorff-Larsen, Improved predictions of phase behaviour of intrinsically disordered proteins by tuning the interaction range, Open Research Europe 2 (2022).

[16] G. Tesei, A. I. Trolle, N. Jonsson, J. Betz, F. E. Knudsen, F. Pesce, K. E. Johansson, and K. Lindorff-Larsen, Conformational ensembles of the human intrinsically disordered proteome, Nature 626, 897 (2024).

[17] R. M. Regy, G. L. Dignon, W. Zheng, Y. C. Kim, and J. Mittal, Sequence dependent phase separation of protein-polynucleotide mixtures elucidated using molecular simulations, Nucleic acids research 48, 12593 (2020).

[18] Z. Benayad, S. von Bulow, L. S. Stelzl, and G. Hummer, Simulation of fus protein condensates with an adapted coarse-grained model, Journal of chemical theory and computation 17, 525 (2020).

[19] J. A. Joseph, A. Reinhardt, A. Aguirre, P. Y. Chew, K. O. Russell, J. R. Espinosa, A. Garaizar, and R. Collepardo-Guevara, Physics-driven coarse-grained model for biomolecular phase separation with near-quantitative accuracy, Nature Computational Science 1, 732 (2021).

[20] Weber, D. Zwicker, F. Jülicher, and C. F. Lee, Physics of active emulsions, Reports on Progress in Physics 82, 064601 (2019).

[21] J. Berry, C. P. Brangwynne, and M. Haataja, Physical principles of intracellular organization via active and passive phase transitions, Reports on Progress in Physics 81, 046601 (2018).

[22] J. Macia and R. V. Solé, Synthetic turing protocells: vesicle self-reproduction through symmetry-breaking instabilities, Philosophical Transactions of the Royal Society B: Biological Sciences 362, 1821 (2007).

[23] R. Wittkowski, A. Tiribocchi, J. Stenhammar, R. J. Allen, D. Marenduzzo, and M. E. Cates, Scalar φ 4 field theory for active-particle phase separation, Nature communications 5, 4351 (2014).

[24] T. A. Witten Jr and L. M. Sander, Diffusion-limited aggregation, a kinetic critical phenomenon, Physical review letters 47, 1400 (1981).

[25] J. J. Christensen, K. Elder, and H. C. Fogedby, Phase segregation dynamics of a chemically reactive binary mixture, Physical Review E 54, R2212 (1996).

[26] S. C. Weber and C. P. Brangwynne, Inverse size scaling of the nucleolus by a concentration-dependent phase transition, Current Biology 25, 641 (2015).

[27] W. M. Jacobs and D. Frenkel, Phase transitions in biological systems with many components, Biophysical journal 112, 683 (2017).

[28] R. P. Sear and J. A. Cuesta, Instabilities in complex mixtures with a large number of components, Physical review letters 91, 245701 (2003).

[29] T. S. Harmon, A. S. Holehouse, and R. V. Pappu, To mix, or to demix, that is the question, Biophysical journal 112, 565 (2017).

[30] Weber, C. F. Lee, and F. Jülicher, Droplet ripening in concentration gradients, New Journal of Physics 19, 053021 (2017).

[31] V. A. Volpert, Elliptic partial differential equations, Vol. 1 (Springer, 2011).

[32] D. Maltseva, S. Chatterjee, C.-C. Yu, M. Brzezinski, Y. Nagata, G. Gonella, A. C. Murthy, J. C. Stachowiak, N. L. Fawzi, S. H. Parekh, et al., Fibril formation and ordering of disordered fus lc driven by hydrophobic interactions, Nature chemistry 15, 1146 (2023).

