## Supplemental figures and methods for "A reaction-diffusion model captures the essence of liquid-liquid phase separation"

### Supplemental Material: Rethinking of droplet formation through liquid-liquid phase separation using reaction-diffusion framework


Nayana Mukherjee, Abdul Wasim, and Jagannath Mondal*

*Tata Institute of Fundamental Research Hyderabad, Telangana 500046, India*

Pushpita Ghosh*

*School of Chemistry, Indian Institute of Science Education and Research, Thiruvananthapuram, Kerala 695551, India*


##### A. Calculation of the free energies using MSD

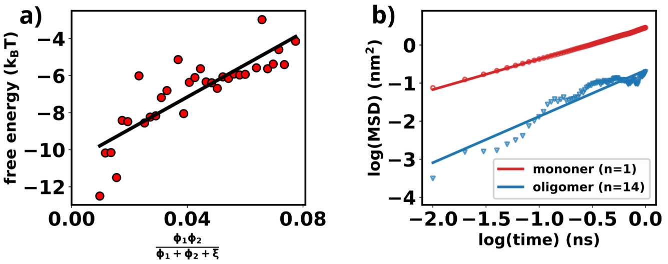


FIG. 1: (a) Free energy vs. $\frac{\phi_i\phi_j}{\phi_i+\phi_j+\xi}$ plot for $\alpha$-synuclein in water with 50 mM of NaCl. (b) Comparison of MSD for dense and dilute phases.

We consider a distribution of $\alpha$-synuclein in various stages of aggregation, where the mole fraction $(\frac{n_i}{N})$ of each cluster or monomer is denoted as $\phi_i$. The parameter '$\xi$' represents the minimum concentration of the dilute phase $P$. Thus, $\phi_i$ ranges from 0.02, signifying a single chain, to 1, indicating complete aggregation of all monomers into a giant cluster. For each pair of $(\phi_i, \phi_j)$, we calculate their closest distance of approach throughout the simulation trajectory. Let $P(d_{ij})$ represent the distribution of a pair of mole fractions $(\phi_i, \phi_j)$. The free energy for such a pair is defined as $min[-\ln(P(d_{ij}))]$, after shifting $P(d_{ij})$ in the y-axis to ensure $max[P(d_{ij})] = 0$. The calculation of free energy $(F)$ values is performed for all observable sets of $(\phi_i, \phi_j)$ during molecular simulations. These $F$ values are plotted on the y-axis against their corresponding $\frac{\phi_i\phi_j}{\phi_i+\phi_j+\xi}$ in Figure 1(a). The Pearson correlation coefficient between the values of $F$ and $\frac{\phi_i\phi_j}{\phi_i+\phi_j+\xi}$ offers insights into the degree of alignment in their trends [1].

##### B. Calculation of the diffusivities of the phases using MSD

Coarse grained molecular dynamics simulations using a modified Martini 3, adapted for $\alpha - synuclein$ [2], were performed on an aggregate (14 mer) and a single chain of $\alpha$-synuclein for determination of their mean squared displacements (MSDs). An initial equilibration in a NPT ensemble for 100 ns with a timestep of 20 ps was performed with the temperature maintained at 310.15 K and the pressure at 1 bar using the v-rescale and the c-rescale thermostats and barostat respectively. This was followed by 2 ns of simulation for each case using a timestep of 0.0001 fs with a data writing frequency of 1000 steps generating 200 frames for each 2 ns trajectory. MSD was then calculated using these trajectories. We got the ratio of $D_{monomer}$ to $D_{oligomer}$ to be 12.52 according to the simulations. So we use the ratio for the droplet to dilute phase to be 1/10.

##### C. Stability analysis of the model

Here we perform linear stability analysis in detail of the system of dimensionless equations:

$$\frac{\partial N(x,y,t)}{\partial t} = \Delta(d_1 N + d_a NP) + N(1 - J(N)) - \frac{aNP}{N+P+\xi} \tag{1a}$$

$$\frac{\partial P(x,y,t)}{\partial t} = \Delta(d_2 P + d_b NP) + \frac{bNP}{N+P+\xi} - cP, \tag{1b}$$

$$\frac{\partial W(x,y,t)}{\partial t} = d_c \Delta W - N(1 - J(N)) + cP + \frac{(a-b)NP}{N+P+\xi}. \tag{1c}$$

where

$$\Delta \equiv \frac{\partial^2}{\partial x^2} + \frac{\partial^2}{\partial y^2}; \quad J(N) = \frac{1}{L^2}\int_0^L \int_0^L N dx dy$$

. Firstly we examine the stability of the temporal dynamics from the temporal model given by (2):

$$\frac{dN}{dt} = N(1-N) - \frac{aNP}{N+P+\xi} \tag{2a}$$

$$\frac{dP}{dt} = \frac{bNP}{N+P+\xi} - cP, \tag{2b}$$

$$\frac{dW}{dt} = -N(1-N) + cP + \frac{(a-b)NP}{N+P+\xi}. \tag{2c}$$

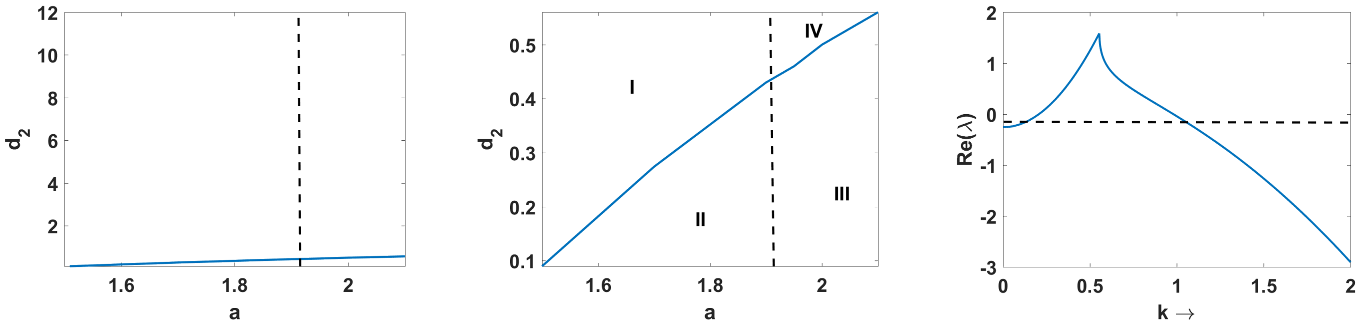


(a) (b) (c)

FIG. 2: (a) Two parameter bifurcation diagram in $a - d_2$ plane for parameter values: $b = 1.0$, $c = 0.6$, $\xi = 0.001$, $d_1 = 1.0$, $d_3 = 10 = d_a = d_b$, black dashed straight line represents the Hopf bifurcation curve, blue curve represents the diffusion induced instability curve, red '$*$' represents the parameter values $a = 1.9, d_2 = 10$ which is used in numerical simulations; (b) Zoomed version of plot in (a). (c) Plot of $Re(\lambda)$ versus wavenumber $k$ for parameter values: $a = 1.9$, $b = 1.0$, $c = 0.6$, $\xi = 0.001$ and $d_1 = 1.0$, $d_2 = 10 = d_3 = d_a = d_b$.

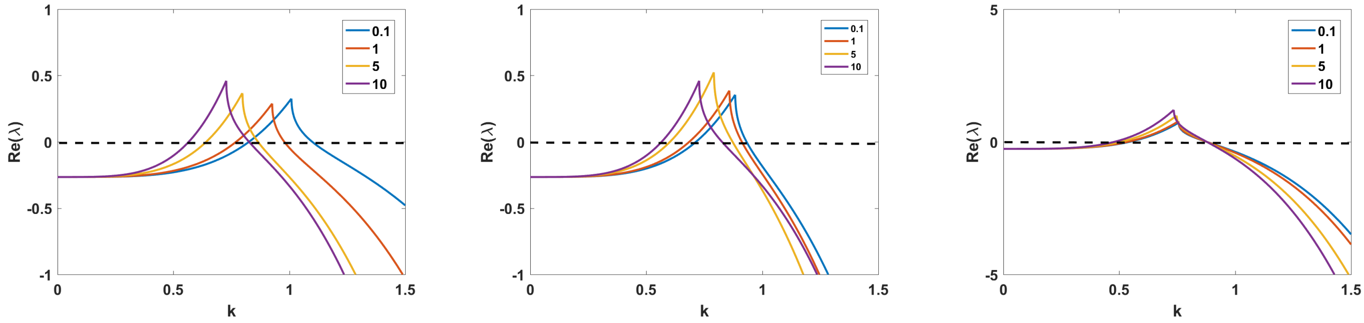


(a) (b) (c)

FIG. 3: Plots of $Re(\lambda)$ over wavenumber $k$ for parameter values are taken as $b = 1$, $c = 0.6$, $\xi = 0.001$ $d_1 = 1$ and varied values of the cross-diffusion parameters $d_a = d_b = 0.1, 1, 5, 10$ (a)$d_2 = d_3 = 0.1$ ;(b) $d_2 = d_3 = 1$ ; (c) $d_2 = d_3 = 5$.

The homogeneous steady state of the system is given by $(N_*, P_*, W_*)$ which are obtained by setting $\dot{N} = \dot{P} = \dot{W} = 0$. We perturb the system around the steady state as by infinitesimal small disturbances: $= N_* + u_1(t)$, $= P_* + v_1(t)$ and $w = W_* + w_1(t)$ where $|u_1|,|v_1|,|w_1| \ll 1$ and obtain the linearized system as follows:

$$\frac{d}{dt}\begin{pmatrix} u_1 \\ v_1 \\ w_1 \end{pmatrix} = \begin{pmatrix} F_1 & F_2 & 0 \\ G_1 & G_2 & 0 \\ H_1 & H_2 & 0 \end{pmatrix} \begin{pmatrix} u_1 \\ v_1 \\ w_1 \end{pmatrix}$$

where $F_1 = (1 - 2N_*) - \frac{aP_*^2 + akP_*}{(N_*+P_*+k)^2}$,
$F_2 = -\frac{aN_*^2 + akN_*}{(N_*+P_*+k)^2}$,
$G_1 = \frac{bkP_* + bP_*^2}{(N_*+P_*+k)^2}$,
$G_2 = \frac{bN_*^2 + bkN_*}{(N_*+P_*)^2} - c$,
$H_1 = -1 + 2N_* + \frac{(a-b)(kP_* + P_*^2)}{(N_*+P_*+k)^2}$
$H_2 = c - \frac{(a-b)(kN_* + N_*^2)}{(N_*+P_*+k)^2}$.

Conditions for local asymptotic stability of $(N_*, P_*, W_*)$ is given by the characteristic equation

$$\lambda(\lambda^2 - \lambda(F_1 + G_2) + F_1G_2 - F_2G_1) = 0$$

We determine the eigenvalues, $\lambda$ from the equation which are 0, $\frac{(F_1+G_2)}{2} \pm \frac{(F_1G_2 - F_2G_2)^{\frac{1}{2}}}{2}$. Depending on the eigenvalues, the equilibrium point is stable for $Re(\lambda < 0)$ but in this case one eigenvalue is always zero. But we can find the Hopf-bifurcation threshold for which the limit cycles or periodic solutions can be seen from the trace of the Jacobian i.e $F_1 + G_2 = 0$.

Now we consider the spatial model (1) without the $W$ part as well as the global average term. That converts the model with non-negative initial conditions and no-flux boundary conditions as into a simplified form as:

$$\frac{\partial N(x,y,t)}{\partial t} = N(1-N) - \frac{aNP}{N+P+k} + \Delta(d_1N + d_aNP) \quad (3a)$$

$$\frac{\partial P(x,y,t)}{\partial t} = \frac{bNP}{N+P+k} - cP + \Delta(d_2P + d_bNP) \quad (3b)$$

Obviously, $N(x,y,t) = N_*$, $P(x,y,t) = P_*$, where $(N_*, P_*)$ is the coexisting equilibrium point of the corresponding temporal model, satisfy system (3) and the associated boundary conditions. Hence we consider $N(x,y,t) \equiv N_*$, $P(x,y,t) \equiv P_*$ to be the homogeneous steady state for (3). Diffusion driven instability occurs when the homogeneous steady-state becomes unstable due to small amplitude heterogeneous perturbations around the homogeneous steady-state. Linear stability analysis for the spatio-temporal model (3) around the homogeneous steady state $E_*(N_*, P_*)$ leads to the conditions for the diffusive instability. We apply small perturbation to the homogeneous steady state as

$$N = N_* + \epsilon_1 \exp((k_x x + k_y y)i + \lambda t) \quad (4a)$$
$$P = P_* + \epsilon_2 \exp((k_x x + k_y y)i + \lambda t) \quad (4b)$$

where $\lambda$ is the growth rate of perturbations and $0 < \epsilon_1, \epsilon_2 \ll 1$. Also, $\mathbf{k} = (k_x, k_y)$ is the wave number vector and $k = |\mathbf{k}|$ is the wave number. After substituting (4) into (3), the characteristic equation for the growth rate $\lambda$ is found from $\mathrm{Det}(\mathbf{J}_1) = 0$, where

$$\mathbf{J}_1 = \begin{pmatrix} F_1 - (d_1 + d_a P_*)k^2 - \lambda & F_2 - d_a N_* k^2 \\ G_1 - d_b P_* k^2 & G_2 - (d_2 + d_b N_*)k^2 - \lambda \end{pmatrix}. \quad (5)$$

By determining the eigenvalues we can check the stability of the system around the homogeneous steady-state. Now, coming to the final model (1), the global average term $J(N)$ becomes constant after a point of term so during the linear stability analysis we consider it to be a constant, say $A$. We use the similar procedure and consider spatio-temporal perturbations around the homogeneous steady-state $E_*(N_*, P_*, W_*)$ as in (4) and get the characteristic equation for the growth rate $\lambda$ is found from $\mathrm{Det}(\mathbf{J}_2) = 0$, where

$$\mathbf{J}_2 = \begin{pmatrix} F_1^{'} - (d_1 + d_a P_*)k^2 - \lambda & F_2 - d_a N_* k^2 & 0 \\ G_1 - d_b P_* k^2 & G_2 - (d_2 + d_b N_*)k^2 - \lambda & 0 \\ H_1^{'} & H_2 & -d_3 k^2 - \lambda \end{pmatrix}, \quad (6)$$

where $F_1^{'} = (1 - A) - \frac{aP_*^2 + akP_*}{(N_* + P_* + k)^2}$, $H_1^{'} = -1 + A + \frac{(a-b)(kP_* + P_*^2)}{(N_* + P_* + k)^2}$.

The characteristic equation is given by:

$$(\lambda + d_3 k^2)(\lambda^2 - \lambda(F_1^{'} + G_2 + k^2(d_1 + d_2 + d_a N_* + d_b P_*) + C(k^2)) = 0,$$

where

$$\begin{aligned} C(k^2) &= C_1(k^2)^2 + C_2(k^2) + C_3 \\ C_1 &= d_1 d_2 + d_a d_2 P_* + d_b d_1 N_* \\ C_2 &= d_a N_* G_1 + d_b P_* F_2 - G_2(d_1 + d_a P_*) - F_1^{'}(d_2 + d_b N_*) \\ C_3 &= F_1^{'} G_2 - F_2 G_1. \end{aligned}$$

By putting the parameter values we can calculate the eigenvalues from the characteristic equation, based on which the stability of the homogeneous steady states can be judged.


[1] P. C. Souza, R. Alessandri, J. Barnoud, S. Thallmair, I. Faustino, F. Grünewald, I. Patmanidis, H. Abdizadeh, B. M. Bruininks, T. A. Wassenaar, et al., Nature methods **18**, 382 (2021).

[2] A. Wasim, S. Menon, and J. Mondal, bioRxiv pp. 2023–10 (2023).
